# SPARC is a new driver of early breast tumor progression via TGF-β -dependent mechanism

**DOI:** 10.1101/2025.01.17.632337

**Authors:** Marianela Sciacca, María del Pilar Carballo, Ezequiel Lacunza, Lina Marino, Paola Noelia Cardozo, Naira Rodríguez Padilla, Tamara López-López, Martin Abba, Laura Courtois, Ivan Bieche, Anne Vincent-Salomon, Érica Rojas Bilbao, Katrina Podsypanina, Mathieu Boissan, Philippe Chavrier, Ana María Eiján, Pablo J Sáez, Catalina Lodillinsky

## Abstract

Ductal carcinoma in situ (DCIS) is a pre-invasive lesion that is thought to be a precursor of invasive ductal carcinoma (IDC). The challenge lies in discriminating between DCIS progressors and DCIS non-progressors, often resulting in over- or under-treatment in many cases. Membrane type 1 (MT1)-matrix metalloproteinase (MMP) has been previously identified as an essential gene involved in DCIS progression. Here, RNA-sequencing analysis of MT1-MMP^high^ subpopulation derived from invasive breast tumors in the intraductal xenograft model was compared against a dataset of human high-grade DCIS, and Secreted Protein Acidic and Cysteine Rich (SPARC) has emerged as a master candidate involved in early breast tumor progression.

We report that SPARC is up-regulated in DCIS as compared to normal breast epithelial tissues, and further increased in IDC relative to synchronous DCIS foci. We found a positive correlation between SPARC and MT1-MMP expression in DCIS lesions. At the mechanistic level, depletion of SPARC reduced MT1-MMP expression, the degradative capacity of the cells and the activation of the TGF-β signalling canonical pathway. Pharmacological inhibition of the TGF-β signalling pathway decreased SPARC and MT1-MMP at the mRNA and protein level, and concomitantly the cell degradative capacity and 3D cell migration. Strikingly, inhibition of the TGF-β signalling pathway limits the invasive transition of breast tumors in a new triple-negative mouse intraductal syngeneic xenograft model. Moreover, high SPARC expression was positively correlated with both, TGF-β and its receptor, TGFBRI, in a basal type of breast cancer collection supporting our findings. This study identifies SPARC as a new driver of early breast tumor progression via a TGF-β-dependent mechanism, suggesting TGF-β signaling pathway as a potential target for patients with high SPARC expression.

## Introduction

Female breast cancer is the most commonly diagnosed cancer type globally as about 2.3 million women are diagnosed with breast cancer, and more than 0.65 million women die from the disease annually^1^. The consequences of this disease highlight the need for new molecular targets to predict recurrence and/or progression and improve existing therapies.

Depending on the absence or presence of invasion, breast carcinomas are classified as DCIS, defined as an intraductal neoplastic proliferation of epithelial cells that is separated from the breast stroma by an intact layer of basement membrane and myoepithelial cells^2^, or invasive carcinoma (IDC). Although 90% of DCIS are asymptomatic, diagnosis of DCIS has increased considerably with the implementation of population-scale screening imaging methods, currently accounting for up to 30% of new diagnosed cancers^3^. DCIS comprises a heterogeneous group of neoplastic lesions that differ in their clinical presentation, histological aspects, as well as genomic, molecular and immune profiles^4–6^. DCIS is generally found adjacent to IDC in the primary tumor. It is considered a (nonobligate) precursor to IDC, but the features that predict progression to more advanced stages are not well defined. Most women with DCIS choose to have breast-sparing surgery. Radiotherapy is indicated in patients undergoing conservative surgery. Hormonal treatment can reduce risk of ipsilateral breast cancer recurrence in women with hormone receptor-positive DCIS^7^. Yet patients with receptor-negative tumors do not have specific treatment. This context shows the importance of identifying new molecular targets for prediction and treatment.

MT1-MMP is overexpressed at the invasive front of invasive breast carcinomas and plays an essential role in the DCIS-to-IDC transition^8^. In the present study, we used transcriptomic data obtained from MT1-MMP^high^ and MT1-MMP^low^ cell population subsets from intraductal MCF10DCIS.com tumor xenografts that identified SPARC as one of the main genes associated with MT1-MMP expression. Here we report an up-regulation of SPARC in DCIS as compared to normal peritumoral breast tissues and further increase in invasive disease components. SPARC expression in advanced stages is not associated with disease progression, identifying SPARC as a gene associated with the early progression of human breast cancer. SPARC expression correlates with MT1-MMP in patient’s samples. In vitro, a reduction in SPARC expression lead to a reduction in MT1-MMP. Furthermore, we found transforming growth factor Beta (TGF-β) highly interconnected with SPARC and pharmacological inhibition of TGF-β receptor type I (TGFBRI) reduces both SPARC and MT1-MMP expression. We also set up a new triple negative murine intraductal syngenic model to investigate how TGFRI signalling pathway influence DCIS-to-IDC transition. Galunisertib decreased the proportion of invasive tumors. Thus, we propose that blockade of the TGFBRI/SPARC/MT1-MMP axis, may offer therapeutic improvements to patients with in situ tumors.

## Materials and Methods

### Cell Culture

MCF10DCIS.com (MCF10DCIS herein) cell line (CVCL_5552) was purchased from Asterand (Detroit, MI, USA) and maintained in advanced DMEM/F12 media supplemented with 10% fetal bovine serum (FBS) and 2 mM glutamine at 37°C in 5% CO_2_. Murine breast cancer cell line LM38-LP (CVCL_B7PY)^9^ was maintained in DMEM-F12 medium (Gibco) supplemented with 2 mM L-glutamine, 80 µg/ml gentamycin and 10% FBS (Internegocios, Argentina) in a humidified atmosphere with 5% CO_2_ at 37°C. Serial passages of confluent monolayers were performed by detaching cells with trypsin (0.25% trypsin and 0.075% EDTA in Ca^2+^ and Mg^2+^ free PBS). All experiments were performed with mycoplasma-free cells and the human cell line, MCF10DCIS has been authenticated using STR profiling.

### Cell Isolation, transcriptomic analysis and Target Gene Selection

Invasive tumors generated after intraductal injection of MCF10DCIS cells in the mammary gland of SCID mice^8^ were harvested and mechanically dissociated. FACS based on human CD298 (BD Pharmingen™, clone LNH-94) and MT1-MMP (Millipore Anti-MMP-14, catalytic domain, clone LEM-2/15.8) antibodies was performed. Based on MT1-MMP expression levels, two different cell populations defined as MT1-MMP**^high^** and MT1-MMP**^low^** were obtained. After RNA purification, samples were processed according to the protocol of SMARTer stranded total RNA seq Kit-Pico Input Mammalian, and then sequenced using HISeqTM, of Illumina at the *Core Facility* of the Institut Curie, Paris, France.

Briefly, FASTQ files generated from sequencing each sample were used to align the sequencing reads to the human genome, producing BAM files. To perform this alignment, we employed the SubRead package within the R/Bioconductor environment. Following alignment, we used Trimmomatic to trim the reads, ensuring data quality and accuracy. The BAM files, after alignment and trimming, were then processed to create a read counts matrix. This matrix served as the input for DESeq2, another R/Bioconductor package, which was utilized to identify differentially expressed genes between the two study groups. The entire analysis was conducted within the R statistical environment

To identify similarities in gene expression between our experimental model and human high-grade DCIS samples, we compared the list of genes upregulated in the MT1-MMP^high^ population of our model with those identified in the human DCIS-C1 and DCIS-C2 subsets. As described by Abba et al. (2015)^10^, these two DCIS subgroups were defined based on their tumor-intrinsic subtypes, proliferative and immune scores, and activity of specific signaling pathways. The more aggressive DCIS-C1 subgroup, which is highly proliferative and basal-like or ERBB2(+), exhibited signatures of activated regulatory T cells and immunosuppressive complexes, reflecting a tumor-associated immune suppression, indicative of a tumor-associated immunosuppressive phenotype. This classification provided a valuable framework for comparing the gene expression profiles of our experimental model with those of human DCIS.

### Human Tissue samples

Formalin-Fixed Paraffin-Embedded (FFPE) human breast tumor tissue were collected from three different cohort of patients. PIC-BIM cohort: 116 primary breast tumor samples were collected at Institut Curie from 2005 to 2006 prior to any radiation, hormonal or chemotherapy treatment. Roffo cohort I: 58 primary breast tumors were collected from 2015 to 2016 and Roffo cohort II: 57 samples of primary breast tumors harbouring IDC and lymph node metastasis were collected from 2015 to 2020 at Instituto de Oncología A. H. Roffo (SI Table S1).

All women provided a signed informed consent for future biomarker research studies. Data were analyzed anonymously. This study complies with the Declaration of Helsinki, and was approved by each institution’s internal review and ethics board (Comité de Pilotage du Groupe Sein, Institut Curie). Analysis of the human samples by immunohistochemistry (IHC) was performed after approval by review board and ethics committee of Instituto Angel H. Roffo (21/18).

Tumor breast molecular subtypes were defined as follows according to the guidelines of the American Society of Clinical Oncology (ASCO)/College of American Pathologists^11,12^. Luminal A: estrogen-receptor (ER)≥10%, progesterone-receptor (PR)≥20%, Ki-67 <14%; Luminal B: ER≥10%, PR<20%, Ki-67≥14%; HER2+: ER<10%, PR<10%, HER2 2+ amplified or 3+; Triple negative breast cancers (TNBC): ER<10%, PR<10%, HER2 0/1+ or 2+ non-amplified. Clinical and pathological features of patients are summarized in Table S1.

### siRNA treatment, Cell viability assay and cell count assay

Small interfering RNA (siRNA) transfection was performed using 5 nM siRNA with Mission®siRNA Trasfection Reagent (Sigma) according to the manufacturer’s instructions. The following siRNAs were used: esiRNA cDNA target sequence Human siSPARC: EHU002941, Mouse siSPARC: EMU088951; MISSION® siRNA Universal Negative Control #1: sic001 (Sigma). Subconfluently monolayers of tumoral cells (LM38-LP or MCF10DCIS) were silenced using siRNA SPARC (according to company specifications). 3x10^3^ cells/100 µl were cultured in 96-well plates and RTq-PCR, IF, gelatin degradation assay and cell viability was analyzed 72 h after transfection.

For testing the involvement of the TGF-β pathway on cell viability, LM38-LP cell line was treated with TGF-β (2 ng/ml, Peprotech) or SB431542 (20 ng/ml, StemCell) for 48 h. Cell viability was determined by the Cristal violet assay (Promega).

### Quantitative real time PCR

Total RNA from murine or human cell lines, were isolated with TRIZol Reagent (Invitrogen) as described^13^. Briefly, cDNA was synthesized using RT-PCR SuperScript™ III One-Step (Invitrogen) and used as template for qPCR analyzing using TransStart Green qPCR SuperMix (TransGen Biotech) and reactions were carried out in the CFX96 Real-Time System, C1000-Thermal-Cycler. Specific primers for mouse, SPARC Forward: 5’- GCCTGGATCTTCTTTCTCCT-3’, Reverse: 5’-GTTTGCAATGATGGTTCTGG-3’; MT1- MMP Forward: 5’-GCTTTACTGCCAGCGTTC-3’, Reverse: 5’- CCCACTTATGGATGAAGCAAT-3’ was normalized to GAPDH (glyceraldehyde-3-phosphate dehydrogenase) expression Forward: 5’-CAAAATGGTGAAGGTCGGTG-3’, Reverse: 5’- CAATGAAGGGGTCGTTGATG-3’. Human primers: SPARC Forward: 5’- GCGGAAAATCCCTGCCAGAA-3’; Reverse: 5’-GGCAGGAAGAGTCGAAGGTC-3’; MT1-MMP Forward: 5’-CAACATTGGAGGAGACACCCACT-3’, Reverse: 5’- CCAGGAAGATGTCATTTCCATTCA-3’ was normalized to TBP (TATA box-binding protein) expression Forward: 5’-TGCACAGGAGCCAAGAGTGAA-3’, Reverse: 5’- CACATCACAGCTCCCCACCA-3’.

For RT-qPCR on the samples of the breast cancer cohort, total RNA extraction, cDNA synthesis and RT-qPCR reaction have been described elsewhere^14^. Quantitative values were obtained from the cycle number (Ct value) using QuantStudio 7 Flex real-time PCR system (Applied Biosystems, Foster City, CA). Data from each sample were normalized on the basis of its content in TBP transcripts. TBP encoding the TATA box-binding protein (a component of the DNA-binding protein complex TFIID) was selected as an endogenous control due to the moderate level of its transcripts and the absence of known TBP retro-pseudogenes (retro-pseudogenes lead to co-amplification of contaminating genomic DNA and thus interfere with RT-PCR transcripts, despite the use of primers in separate exons). The relative mRNA expression level of each gene, expressed as N-fold differences in target gene expression relative to the TBP gene and termed ‘Ntarget’, were determined as Ntarget = 2^ΔCtsample^, where the ΔCt value of the sample was determined by subtracting the average Ct value of target gene from the average Ct value of TBP gene. The Ntarget values of the samples were subsequently normalized such that the median Ntarget value of the normal breast samples was 1. Primers for MT1-MMP (upper primer, 5’- TTGGAGGAGACACCCACTTTGACT-3’; lower primer, 5’- CCAGGAAGATGTCATTTCCATTCAG-3’), SPARC (upper primer, 5’- TGTGGCAGAGGTGACTGAGGTATC -3’; lower primer, 5’- TCGGTTTCCTCTGCACCATCA-3’) and TBP (upper primer, 5’- TGCACAGGAGCCAAGAGTGAA-3’; lower primer, 5’-CACATCACAGCTCCCCACCA-3’), were selected with Oligo 6.0 program (National Biosciences, Plymouth, MN).

Metastasis-free survival (MFS) was determined as the interval between diagnosis and detection of the first metastasis. Survival distributions were estimated by the Kaplan-Meier method, and the significance of differences between survival rates was ascertained using a log-rank test. GraphPad Prism (GraphPad Software) was used for statistical analysis.

### Western blot assay

Subconfluently monolayers of tumor cells (LM38-LP or MCF10DCIS) were silenced for SPARC or were treated with or without SB431542. Then, cells were processed for western blot. Briefly, cells were gently washed with PBS and lysed using protein extraction lysis buffer (50 mM Tris-HCl (pH 8.0); 100 mM NaCl; 1% Triton, 1 mM/ml aprotinin, 1 mM phenylmethylsulfonyl fluoride, 2 mg/ml leupeptin and 10 mM EDTA/EGTA). Protein concentration was determined by Bradford method according to the manufacturer’s instruction (Merk). Aliquots from the cell lysates were separated by electrophoresis and analyzed in 10% sodium dodecyl sulfate-polyacrylamide gel (SDS-PAGE) and transferred onto a PVDF membrane. After blotting, the membrane was incubated with primary antibody followed by a horseradish peroxidase conjugated secondary antibody, for 1 h at room temperature. Blots were developed using the ECL detection kit and ImageQuant LAS 500 CCD imager (GE Life sciences). Membranes were stripped and incubated overnight with Tubulin used as a loading control.

### Immunofluorescence analysis

LM38-LP or MCF10DCIS cells growing in chamber slides with complete medium were silenced for SPARC. Subconfluent monolayers were gently washed with cold PBS and processed for immunofluorescence. Briefly, slides were fixed with PFA 4% in PBS for 15 minutes and permeabilized with Triton X-100 0,3%, with a blocking solution containing 5% FBS in PBS for 1h at room temperature. Fixed cells were incubated overnight with the primary antibody and the following day fixed cells were incubated 2h with secondary antibody. Nuclei were counterstained with DAPI (Santa Cruz Biotechnology) and slides were observed in a Nikon EclipseTM E400 fluorescence microscope and photographed with a CoolpixH 995 digital camera. Mean Fluorescence (AU) was quantified by using Image J software.

### Fluorescent Gelatin Degradation Assay

Assays of fluorescent gelatin degradation were performed and quantified as previously described^15^. Briefly, clean coverslips are incubated with 1:2000 poly-L-lysine (0.5 µg/ml, Sigma) 20 min at room temperature (RT), washed with PBS and incubated with 0.5% glutaraldehyde 15 min RT. Then washed 3 times with PBS and incubated for 10 min at RT the glass slides by inversion on gelatin-A488 (G13186, Invitrogen) placed on Parafilm previously cleaned with 70% ethanol. The coverslip was washed with PBS and incubated with 5 mg/ml sodium borohydrate (Sigma) during 3 min. The coverslip is washed 3 times with PBS, complete medium is added and incubated with cell suspension. The seeded cells were previously treated with TGF-β (2 ng/ml, Peprotech), SB431542 (20 ng/ml, StemCell) or Galunisertib (1 µg/ml) overnight. After 3h of seeding, cells continued to receive the corresponding treatment. Then, cells were fixed 20 min RT with PFA 4% and stained with phalloidin (A22283, Life technologies) and DAPI (Santa Cruz Biotechnology). Slides were observed in a Nikon EclipseTM E400 fluorescence microscope and photographed with a CoolpixH 995 digital camera. The degradation capacity of the cells was determined using Image J, where levels of proteolysis were quantified as negative fluorescent areas.

### Migration assays

To monitor cell migration in 3D complex microenvironments we used collagen gels following an established method^16–18^. The device was fabricated by curing polydimethylsiloxane (PDMS) on a silicone mold with a positive relief of three compartments previously described^17^. The resultant PDMS channel was bonded to a glass bottom Petri dish (Fluorodish) at 65°C for 20 min. Then, rat tail type I collagen (Corning) was diluted to a final concentration of 2 mg/mL with 10% 10X PBS by volume and deionized water, the mixture was prepared and kept at 4°C. The pH was adjusted to pH 8.0 with 1 N sodium hydroxide (NaOH). The cells were added in medium with a final concentration of 0,6x10^6^ cells/ml. The gels polymerized after 20 min of incubation at 37°C. After the administration of the corresponding treatments cellular behavior was monitored using an inverted microscope (Leica Dmi8) equipped with an APO 10x/0.45 PH1, FL L 20x/0.40 CORR PH1, and APO 40x/0.95 objectives. Images were recorded with an ORCA-Flash4.0 Digital camera (Hamamatsu Photonics) using the MetaMorph Version 7.10.3.279 software (Molecular Device). Images were captured every 3 min for 16 h, employing 10x magnification with PH1 phase contrast and binning set to 2.

For 2D migration experiments, 1.5x10^3^ cells were plated on a glass bottom dish and treated for 2 h with TGF-β (2 ng/ml, Peprotech) or SB431542 (20 ng/ml, StemCell) and stained with Hoesch (Invitrogen). Cells were monitored using the same system described for 3D, using 10x magnification and for a total duration of 4.5 h. Cell tracking was performed by combining previously described methods^17,18^. Briefly, raw movies were processed, and later on tracked by using TrackMate v7 plugin from Fiji (ImageJ) to obtain the main migration parameters^17,18^.

### Intraductal tumor growth

Ten-week-old female BALB/c mice obtained from our Institute Animal Care Division were inoculated intraductally in the fourth pair of mammary glands with 5x10^3^ LM38-LP cells as previously described^8,19^. After three weeks mice were randomized into 2 groups: Control and Galunisertib (n=10 per group). After the detection of any palpable tumor (around the fifth week) Galunisertib was orally administered once a day at a dose of 60mg/kg for 5 days. Then, mammary glands were harvested and processed for histological procedures. Mice were handled in accordance with the international procedure for Care and Use of Laboratory Animals. Protocols were approved by the Institutional Review Board CICUAL, Institute of Oncology Angel H. Roffo (04/2023).

### Histological and immunofluorescence analysis of mouse tissue sections

Whole-mount carmine staining and hematoxylin and eosin (H&E) staining of tissue sections were carried out as previously described^20^. After whole mount staining, image acquisition was performed with a Zeiss Stemi 2000-C Stereo Microscope. Quantification of the tumor area was performed using Image J software. To retrieve antigens on paraffin-embedded tissue samples, sections were incubated for 20 min in 10 mM sodium citrate buffer, pH 6.0 at 90°C. Then, after 60 min incubation in 5% FCS, sections were incubated overnight with diluted primary antibodies, washed and further incubated for 2 h at room temperature with appropriate secondary antibodies. Conventional Hematoxylin (Biopur) and Eosin (Merk) staining was carried out according to manufacturer’s instructions.

### Antibodies

Antibodies for immunofluorescence and IHC (conventional or Immunofluorescence) analysis on tissue sections, the following primary antibodies were used: anti-human MT1-MMP (clone LEM-2/15.8, Millipore MAB3328, 1/100) anti-mouse MT1-MMP (PA5-13183, Invitrogen, 1/500), Ki-67 (clone MIB-1, Dako, 1/100), anti-human SPARC (GTX133747 GeneTex), anti-mouse SPARC (PA-5-80062, Invitrogen, 1/500), anti-SMAD-4 (sc-7966, Santa Cruz, 1/200), anti-ki67 (ab1558, Abcam, 1/500), anti-Progesterone receptor A/B (#8757, Cell Signaling, 1/100), anti-Estrogen receptor alfa (sc-542, Santa Cruz, 1/100), anti-HER2/neu (Clon, 4B5, VENTANA anti-HER2/neu, Roche). For immunofluorescence staining of cells in culture: F-actin was stained with Alexa546-phalloidin (A22283, Life technologies, 1/1000) and nuclei with 4’,6-diamidino-2-phenylindole (sc-3598, Santa Cruz, 1 μg/ml). Secondary antibodies Goat anti-rabbit A488 (ab150077, abcam, 1/500) or anti mouse A488 (ab150113, abcam, 1/500) were used. For immunoblotting analysis, we used anti p-SMAD 2/3 (Cell Signalling, D27F4 1/500) anti-SMAD 2/3 (sc-133098, Santa Cruz, 1/100), anti-MT1-MMP (clone MAB3328, Millipore, 1/500), β-tubulin (#2146, Cell Signalling, 1/3000) and detection was performed with anti-rabbit IgG Mouse-HRP (RD#HAF007, R&D Systems, 1/5000) or anti-mouse IgG Rabbit-HRP (RD#HAF008, R&D Systems, 1/5000).

### Gene Expression Data Analysis

To examine the association and co-expression of genes in breast cancer tissues, we employed scatter plots with regression lines. Data for this analysis was sourced from the TCGA PanCancer Atlas. For assessing the expression of SPARC and MT1-MMP in breast cancer cell lines, we utilized RNA sequencing data from the GSE48213 dataset, which includes comprehensive transcriptional profiling of a breast cancer cell line panel. Additionally, timeline data documenting the progression from normal breast tissue to invasive breast cancer was generously provided by Rebbeck et al^21^. To define the network connectivity and functional associations between coexpressed genes, we utilized the STRING web tool (https://string-db.org/), which provides a comprehensive analysis of known and predicted protein-protein interactions. In addition, we employed the Gene Ontology (GO) analysis tools integrated within STRING to further investigate the biological processes, molecular functions, and cellular components associated with these genes, allowing for a deeper understanding of their potential roles in the studied context.

### Statistical analysis

All experiments were repeated almost two times independently. Data are expressed as the mean ± SD or ± SEM. Statistical analyses of H-score, protein and mRNA and protein levels, gelatin degradation assay, cell survival, migration and invasion assays were performed using X^2^ test, Student’s t-test, one-way or two-way analysis of variance using GraphPad Prism (GraphPad Software, La Jolla, CA, USA) as specified in each figure legend with p<0.05 considered significant. For survival analysis, Kaplan–Meier plots were drawn and statistical differences evaluated using the log-rank test. For correlation analysis in cell lines, Pearson’s correlation was used. For boxplots indicating gene expression Wilcoxon rank sum test was used. (R software version 3.0.2, R Foundation for Statistical Computing, Institute for Statistics and Mathematics, Vienna, Austria).

## Results

### Screening for factors involved in the early progression of breast cancer

We previously reported that MT1-MMP expression is up-regulated in invasive versus *in situ* breast carcinomas and that high MT1-MP levels are correlated with poor clinical outcome in breast cancer patients^8^. In addition, we found that MT1-MMP expression is required for the DCIS-to-invasive transition of breast tumor xenografts formed upon intraductal injection of MCF10DCIS cells in the mammary gland of immunocompromised SCID mice^8^. With the aim of identifying new genes that may contribute to the DCIS-to-IDC transition, we performed RNAseq analysis on MT1-MMP^high^ and MT1-MMP^low^ MCF10DCIS cell populations obtained from MCF10DCIS intraductal tumor xenografts.

Comparison of the two groups identified 47 differentially expressed genes (p value<0.05, DR<0.05), of which 46 were up-regulated and one was down-regulated. Computational analysis of the gene list revealed significant enrichment (p<0.01) in biological processes associated with cell adhesion (mediated by integrins) and the extracellular matrix (among others, collagens and laminins). These results are consistent with the canonical role of MT1-MMP in extracellular matrix remodelling, during cell invasion (Figure 1A and Supp file 1).

**Figure 1.**
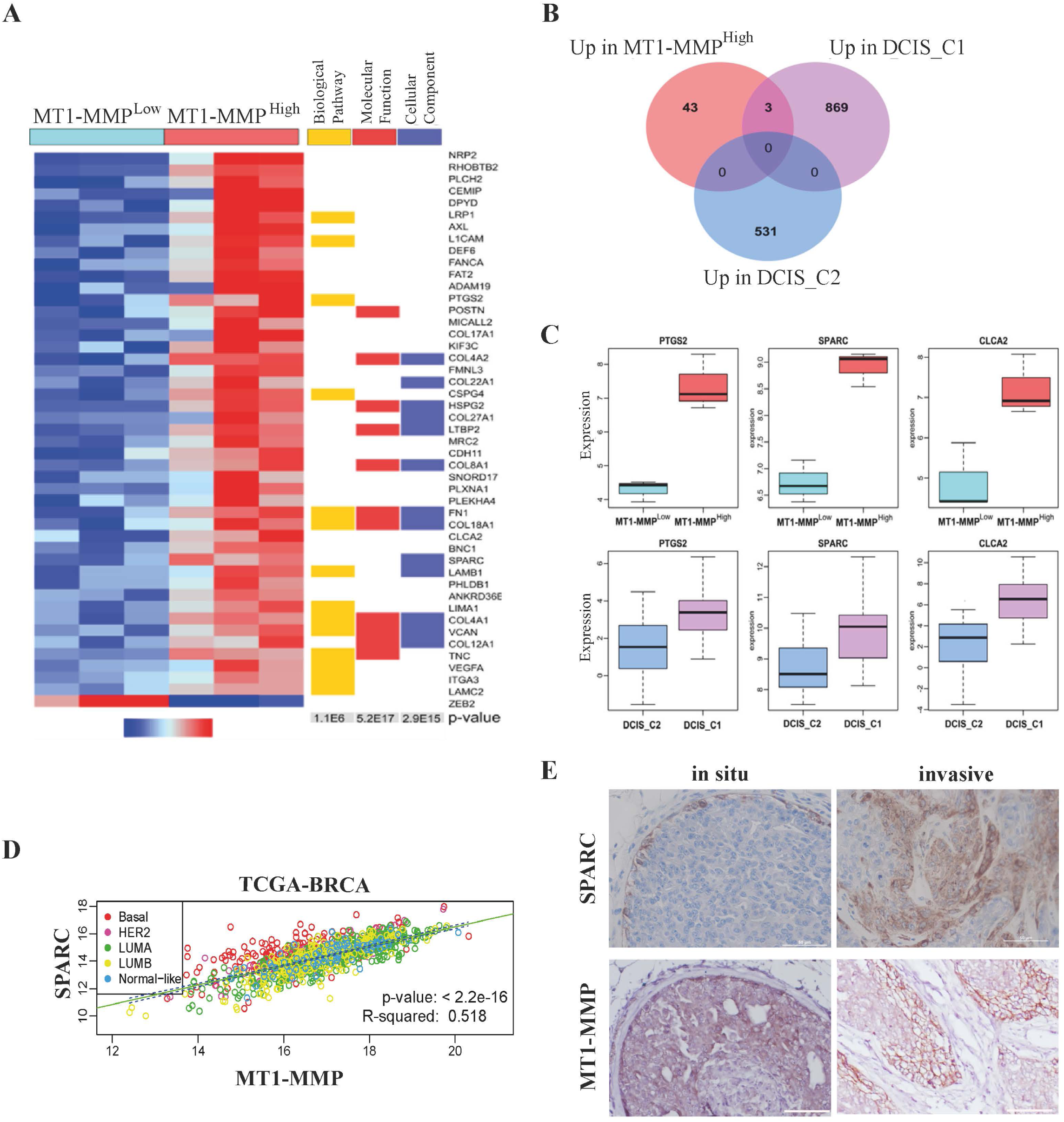
Screening of factors potentially involved in the early progression of breast cancer. **A.** Heat-map obtained after transcriptomic analysis of MT1-MMP^high^ versus MT1-MMP^low^ populations. **B**. Venn diagram of the list of differentially expressed genes between the DCIS-C1 and -C2 subgroups^10^ with the genes upregulated in the MT1-MMP^high^ cell subpopulation. **C**. Box and whisker plots showing the expression of genes in MT1-MMP^high^ and MT1-MMP^low^ in the MFC10DCIS.com cell population and in DCIS_C2 versus C1 groups. Upper panel PTGS2: p-value = 0.01965; SPARC: p-value = 0.0255; CLCA2: p-value = 0.001893 comparing MT1-MMP^high^ versus MT1-MMP^low^. Lower panel: PTGS2: p-value **=** 0.01389; SPARC: p-value **=** 0.002101; CLCA2: p-value = 0.02451. **D**. **S**catter plot displaying the relationship between SPARC and MT1-MMP expression levels in breast cancer tumors classified into the intrinsic subtypes, based on data from the TCGA breast cancer dataset. The plot includes a linear regression line (green) representing the best-fit model between the two variables, along with a 95% confidence interval for the regression predictions (blue dashed lines). The regression analysis shows a correlation coefficient (R) of 0.51 and a p-value < 0.001, indicating a significant positive association between SPARC and MMP14 gene expression in breast cancer tumors. **E**. IHQ illustrating SPARC and MT1-MMP expression in in situ and invasive tumors after intraductal injection of MCF10DCIS. Scale bar: 50 µm.

Abba and colleagues^10^, described two distinct DCIS subgroups (aggressive DCIS-C1 and indolent DCIS-C2) based on tumor-intrinsic subtypes, proliferative,immune scores, and activity of specific signaling pathways. With the objective of finding genes potentially involved in the process of DCIS-to-IDC transition we compared upregulated genes in the two DCIS subgroups—relative to normal breast tissue—with upregulated genes in MCF10DCIS MT1-MMP^high^ cells of the xenograft model. Cross-species comparison identified no overlap with DCIS-C2 signature; however, three genes could be detected both in MT1-MMP^high^ and the high-risk human DCIS-C1 signatures (Figure 1B). These genes, Secreted Protein Acidic and rich in cysteine (SPARC), Cyclooxygenase-2 (PTGS2), and Calcium-Activated Chloride Channel regulator 2 (CLCA2) showed similar relative expression changes in xenografted MT1-MMP^low^ vs. ^high^ cells and in primary DCIS_C1 vs. _C2 subgoups (Figure 1C).

We next examined the correlation between the expression of the three candidate genes and MT1-MMP using public databases. SPARC showed the best correlation with MT1-MMP in breast cancer samples (from TCGA, p-value: < 2.2e-16) (Figure 1D). In situ immunohistochemistry (IHC) on xenograft tumor sections showed that SPARC was overexpressed at the invasive edge of MCF10DCIS tumors, similar to MT1-MMP expression (Figure 1E). Taken together these results indicated that SPARC is a candidate gene potentially involved in the regulation of the invasive switch of breast cancer.

### SPARC expression increases during early breast cancer progression

We hypothesized that breast cancer progression is associated with an increase in SPARC expression. We stained breast tissues including normal, DCIS and IDC components from two different cohorts of patients (PIC-BIM n=78, and Roffo n=34) for SPARC. In line with previous reports^22^, SPARC signal was detected in myoepithelial cells of normal mammary ducts, in tumor-associated fibroblasts and in endothelial cells (data not shown). We focused our analysis on carcinoma cells, in which we observed granular cytoplasmic SPARC staining, and calculated an histo (H)-score value based on the percentage of positive cells multiplied by intensity staining on a 0-3 scale (Figure 2A). Overall, SPARC was undetectable in normal breast epithelial cells, but significantly upregulated in tumor tissue. Moreover, in matched DCIS-IDC samples from both cohorts, SPARC levels were higher in IDC than in synchronous DCIS foci (Figure 2A-C). In addition, the proportion of SPARC-positive IDC tumors was higher in high-grade than in lower-grade breast cancers (Figure 2D). When IDC tumors were stratified into breast molecular subtypes, SPARC expression was higher in TNBC (Figure 2E).

**Figure 2.**
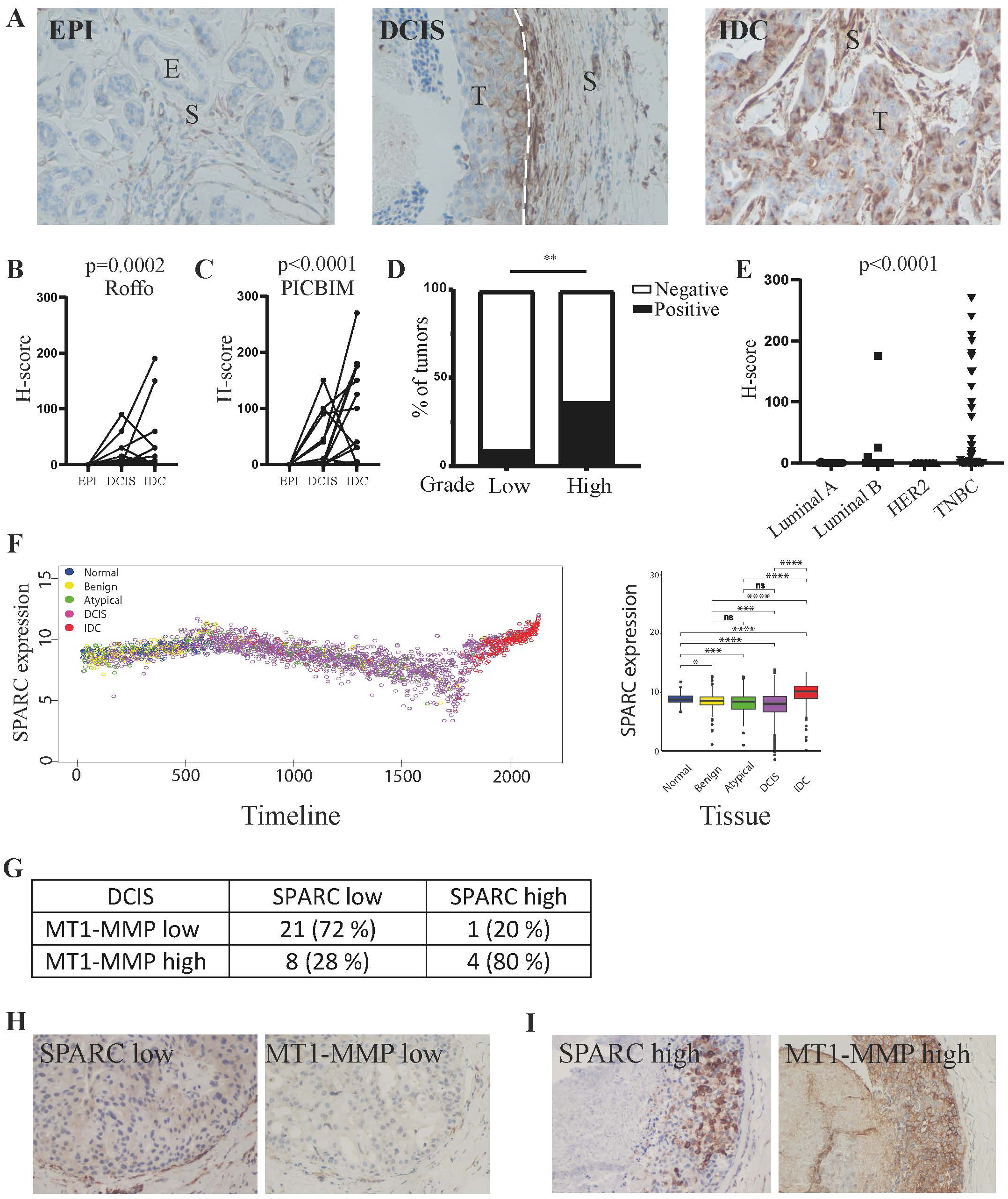
Overexpression of SPARC during human breast cancer progression. **A.** Representative SPARC IHC staining in breast peritumoral tissues and synchronous in situ and invasive components from one breast carcinoma biopsies. Dotted line separates stroma from de DCIS. E: Epithelium; T: Tumor; S: Stroma. Scale bar: 25μm. **B, C.** SPARC levels using the H-score method in the adjacent peritumoral tissues, in situ and invasive breast carcinomas in PICBIM’s cohort (B)***p=0.0002, Friedman test and in Roffo’s cohort (C) ***p<0.0001, Friedman test **D**. Proportion of cases regarding low (I and II) or high (III) histological grade **p=0.002 X^2^ test, two-side. **E**. H-score of SPARC in IDC tumors of Roffo’s cohort regarding molecular subtype, ***p<0.0001 Kruskal-Wallis test. **F**. Left panel: Scatter plot illustrating the trend of SPARC expression -log2(CPM)-versus position along the timeline, during the progression from normal breast tissue to invasive breast cancer. Right panel: Box-plot illustrating SPARC expression into the different groups considered^21^ *p<0.05; ***p<0.001; ****p<0.0001. ns non-significant Wilcoxon rank sum test **G**. Proportion of cases with high or low expression level of SPARC and MT1-MMP of consecutive slides. *p=0.0235 X^2^ test, two-side. **H, I**. Representative IHQ staining in DCIS components for SPARC and MT1-MMP low (H) and high (I) expression levels. Scale bar: 50μm.

Recently, it was proposed that the progression from in situ lesions to IDC involves dual, early and late epithelial-to-mesenchymal transition (EMT) events^21^. Rebbeck and colleagues conducted a pseudo-time analysis using a fitted principal curve on a principal component analysis (PCA) plot of samples, based on the most significant differentially expressed genes between DCIS and co-occurring IDC^21^. This analysis suggested that DCIS samples exhibit varying relationships to their normal counterparts or invasive counterparts, potentially indicating a continuum of tissue states during disease progression within individual patients. To investigate this further, we examined the evolution of SPARC expression across the pseudo-time sequence, correlating it with tumor progression. We found that, similar to EMT genes, SPARC expression exhibited two peaks, suggesting a dynamic role during the transition from DCIS to IDC (Figure 2F, left panel). Consistent with immunohistochemistry (IHC) data, SPARC transcript levels were highest in IDC samples (Figure 2F, right panel). These findings support the idea that SPARC may serve as a potential biomarker for the transition from DCIS to IDC, with its increased expression linked to tumor progression. In order to study the association between SPARC and MT1-MMP, we analyzed MT1-MMP expression in the Roffo cohort. We focused our analysis on carcinoma cells in which MT1-MMP staining was observed at the plasma membrane or as a granular cytosolic pattern. We found a statistically significant fraction of DCIS with high MT1-MMP expression coinciding with high expression of SPARC (Figure 2G-I). Thus, SPARC and MT1-MMP are coordinately expressed in the experimental xenograft model (Figure 1E) as well as in patient samples.

To rule out that SPARC is a general progression factor we examined a third cohort of patients, where samples contain both IDC foci and metastatic axillary lymph node tissue. Importantly, lymph node metastases did not contain SPARC-positive cells, in contrast to the primary tumor (Figure supp 1A-C). In addition, SPARC expression by neoplastic cell in IDC was not associated with Relapse Free Survival (Figure supp 1D). These results suggest that SPARC is specifically involved in early invasive progression. SPARC expression is present in the healthy stroma^22^. In our cohorts, SPARC expression in the tumor stroma did not show significant differences with respect to the molecular subtypes (data not shown). Furthermore, expression of SPARC and MT1-MMP increased concomitantly in cell lines with basal features (Figure supp 2A), consistent with the idea that co-regulation of these genes may be informative only in the neoplastic compartment. To further deepen our analysis, the prognostic power of SPARC and MT1-MMP co-expression was evaluated by RTqPCR in a retrospective cohort of 458 breast cancer patients in which we previously reported that MT1-MMP up-regulation correlated with shorter metastasis-free survival (MFS) (Figure supp 2B, left panel and Ref ^8^). However, neither SPARC nor combination of SPARC and MT1-MMP expression impacted metastasis-free survival (Figure supp 2B, middle and right panel). This result confirmed that SPARC expression both at mRNA and protein levels was unlikely to contribute to dissemination of breast cancer. Taken together, our findings argued that SPARC was involved exclusively in early progression of breast cancer, a mechanism enhanced by concomitant MT1-MMP up-regulation in the aggressive TNBC subtype.

### MT1-MMP mediates the invasive potential of SPARC

With the aim of strengthening our results in an immunocompetent murine model, the role of SPARC in early breast cancer progression was analyzed by using the mouse mammary cancer cell line LM38-LP^9^, which was inoculated in the ductal system of syngeneic mice (Figure sup 4A). Whole-mount and histology staining revealed *in situ* tumors three weeks post-intraductal injection of LM38-LP cells. A continuous layer of elongated α-SMA positive myoepithelial cells segregating *in situ* tumors from the stroma was observed (Figure supp 3B). During the fifth week, LM38-LP tumors spontaneously progressed to invasive lesions characterized by disrupted stromal collagen organization (Figure supp 3AB). These tumors were negative for ER, PR and HER2/neu (Figure supp 3C). Thus, we concluded that LM38-LP xenograft tumors are an appropriate model to study the molecular events involved in the early transition of triple-negative breast cancer in immunocompetent mice. Both the LM38-LP cell line *in vitro* and tumors generated after intraductal inoculation were positive for MT1-MMP and SPARC with a perinuclear, granular distribution (Figure supp 4A). Knockdown of SPARC by RNAi did not affect cell proliferation or morphology (Figure supp 4B and C). Interestingly, reduction in SPARC expression led to a reduction in MT1-MMP expression in both human MCF10DCIS and murine LM38-LP cells, which correlated with reduction of gelatin degradation capacity (Figure 4CD and supp 4DE). Collectively, these data suggested that SPARC played a pro-invasive role mediated by MT1-MMP.

**Figure 3.**
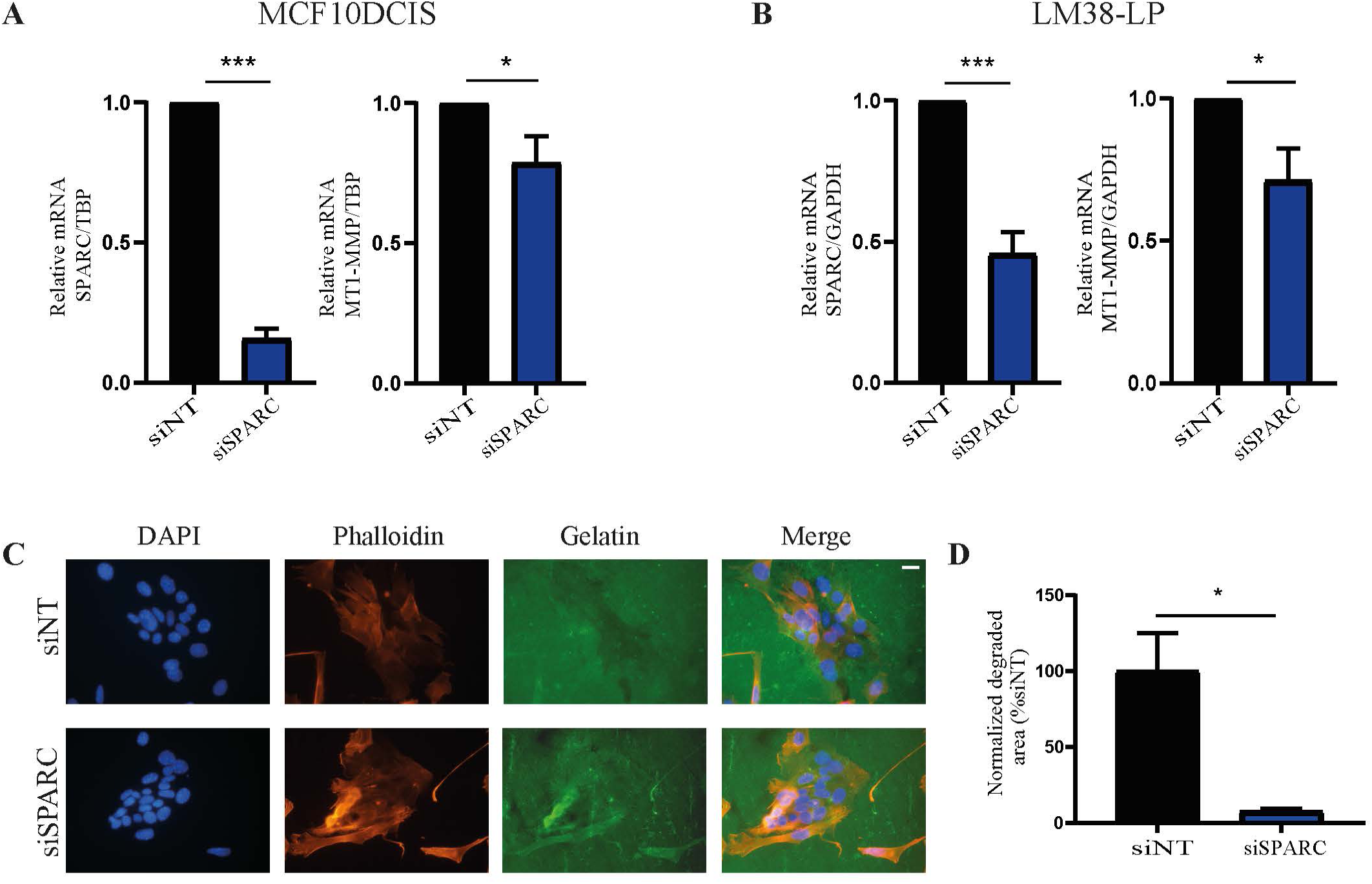
Pro-invasive role of SPARC mediated by MT1-MMP. **A**. LM38-LP murine breast cell line was transiently silenced for SPARC expression. Values corresponding to means ± SEM, n = 3 independent experiments of relative amounts of mRNA normalized against GAPDH and relativized to their CRL. **** p<0.0001, *p<0.05 using one-side t-test. **B.** SPARC and MT1-MMP expression in MCF10DCIS after transiently silenced for SPARC. Values corresponding to means ± SEM, n = 2 independent experiments of relative amounts of mRNA normalized against GAPDH and relativized to their CRL*** p<0.0001, **p<0.01 t-test. **C.** Representative images of cells in gelatin. Blue: DAPI, Green: Gelatin, Red: Phalloidin. Magnification bar: 50µm **D**. Quantification of fluorescent gelatin degradation by total cells in LM38-LP cells silenced or non-silenced for SPARC. Quantification of 1 assay (n=2). *p<0.05 using one-tailed t-test.

**Figure 4.**
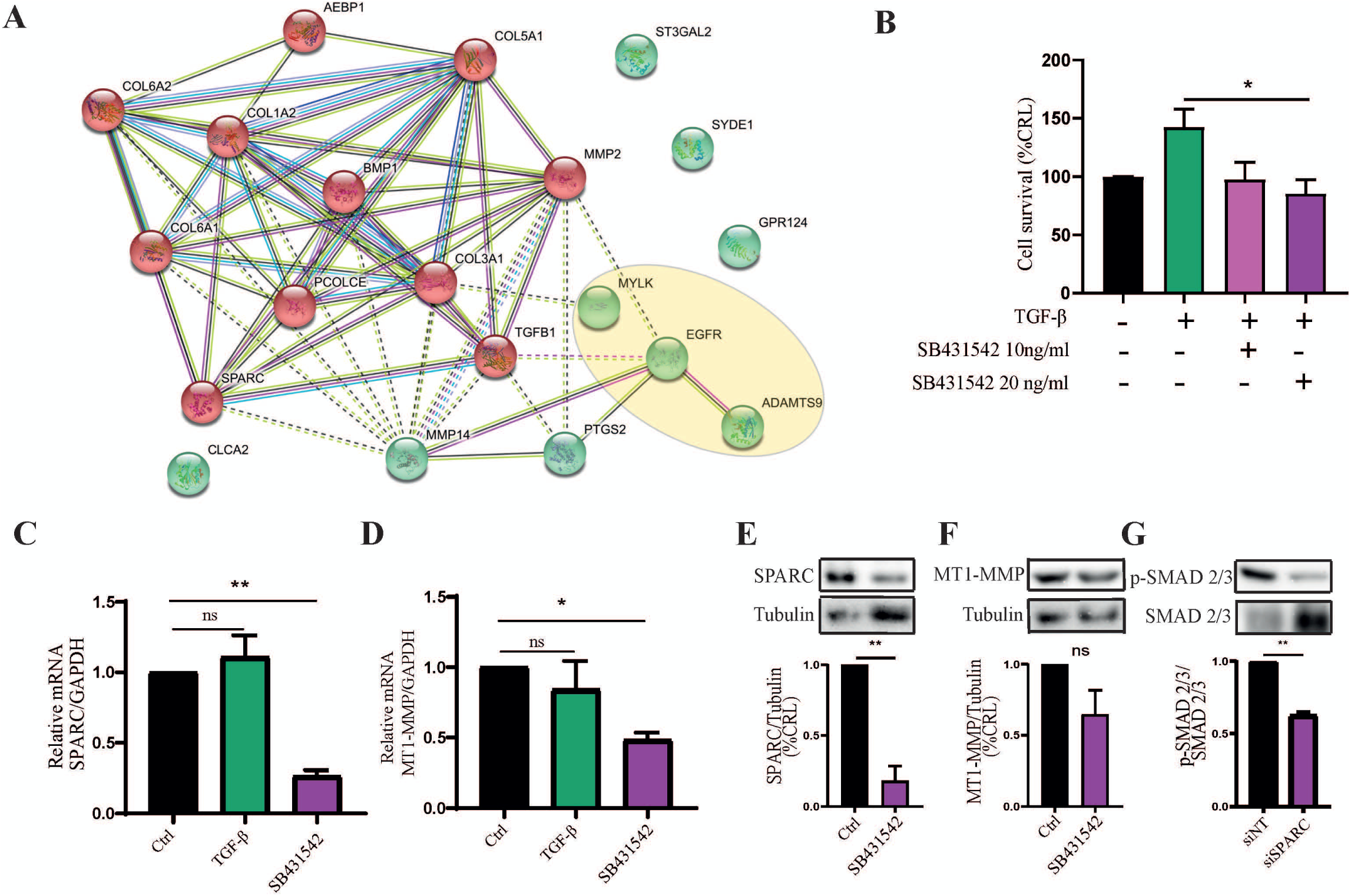
Rol of TGF-β molecular pathway in the SPARC pro-tumoral process. **A.** Association STRING analysis between the four candidate genes and co expressed genes. **B.** LM38-LP cells were treated with TGF-β (2ng/ml) and SB431542 (10 and 20 ng/ml) for 48 h, cell viability was evaluated by crystal violet assay. *p<0.05, **p< 0.01 by two-way ANOVA, Tukey’s multiple comparisons test vs CRL. Values are means ± SEM, n=3 independent experiments. **C, D.** Expression mRNA level of SPARC (C) and MT1-MMP (D) in LM38-LP cells after SB431542 treatment (20ng/ml). Values corresponding to means ± SEM, n = 3 independent experiments of relative amounts of mRNA normalized against GAPDH and relativized to their CRL, t-test, *** p<0.0001. **E, F.** Protein levels of SPARC (E) and MT1-MMP (F) in LM38-LP cells after SB431542 treatment (20 ng/ml). Values corresponding to means ± SEM, n = 3. t-test, **p<0.001. **G.** SMAD 2/3 phosphorylation was determined in LM38-LP cells transiently silenced for SPARC by Western Blot. Representative imaging and quantification. **p=0.0033 Unpaired t test, two-tailed, ns=non-significant.

To profile molecules involved in the early transition process, we used a subset of publicly available breast cancer data from the TCGA database to conducted a gene co-expression analysis of the three previously identified genes, SPARC, PTGS2, and CLCA2 together with MMP14 (encoding MT1-MMP) in five breast cancer sample groups Luminal A (LumA, n=473), Luminal B (LumB, n=205), Her2 (n=86), Normal-like (NL, n=73), and Basal (n=185) tumor subtypes and also in an adjacent normal tissue (NAT, n=103) (Supp file 1).

Analysis of co-expressed genes from each gene list (coefficient (R) > 0.5 and a p-value < 0.0001), irrespective of breast cancer molecular subtypes, revealed three common genes: MYLK, EGFR, and ADAMTS9 co-expressed with all the four candidate genes (Figure supp 5A). SPARC and MMP14 exhibited the highest number of partners (Figure supp 5B). Focusing on genes co-expressed with individual candidate genes across all breast cancer groups, we identified 161 genes for SPARC, 51 for MMP14, 10 for PTGS2 (except in Basal), and none for CLCA2 (Supp file 1). Notably, only SPARC and MMP14 shared common partners (Figure supp 5B): COL6A2, AEBP1, COL5A1, MMP2, BMP1, COL1A2, COL6A1, COL1A2, PCOLCE, COL3A1, ST3GAL2, SYDE1, GPR124. Further analysis of these 13 shared genes and the three genes coexpressed in all five breast cancer groups revealed a strong association among 80% of them (Figure 4A). The network analyses delineate two main clusters whose genes were functionally associated with the Gene ontology terms *collagen, matrix degradation, and the TGF-β signaling pathway* (Figure 4A).

We could experimentally confirm the involvement of the TGF-β pathway in the SPARC-dependent cell survival, as treatment of LM38-LP cells with TGF-β increased cell viability, while treatment with TGFBRI inhibitor, SB431542, abolished the survival response (Figure 4B). Although treatment of LM38-LP cells with TGF-β did not affect SPARC expression (data not shown). Inhibition of TGF-β signaling with SB431542 decreased both SPARC and MT1-MMP expression at the mRNA and protein levels, suggesting that activation of TGFBRI signaling contributes to steady-state SPARC and MT1-MMP expression in both LM38-LP and MCF10DCIS cell lines (Figure 4C-F and supp 5C). To test the involvement of SPARC in canonical activation of the TGF-β pathway, we evaluated the phosphorylation status of SMAD2/3 acting downstream of TGFBRI. In cells knocked down for SPARC, p-SMAD 2/3 levels were reduced (Figure 4G), implying that SPARC is required for optimal activation of the TGF-β signaling pathway. Consistent with the loss of MT1-MMP expression following inhibition of TGFBRI, we observed a decrease in gelatinolysis capacity using two different inhibitors, SB431542 or Galunisertib (Figure 5A-D). To determine the effect of TGF-β pathway modulation on motility, we monitored single cell migration in 3D collagen matrices^17^, which mimic interstitial migration. TGF-β accelerated the mean speed of LM38-LP cells as compared to control, while treatment with SB431542 reduced cell motility in both LM38-LP and MCF10DCIS cell lines (Figure 5E and Figure supp 5D). Interestingly, the effect of SB431542 treatment was lost in silencing of SPARC in LM38-LP cells, confirming that TGFBRI-induced cell motility is at least partially dependent on SPARC (Figure 5F).

**Figure 5.**
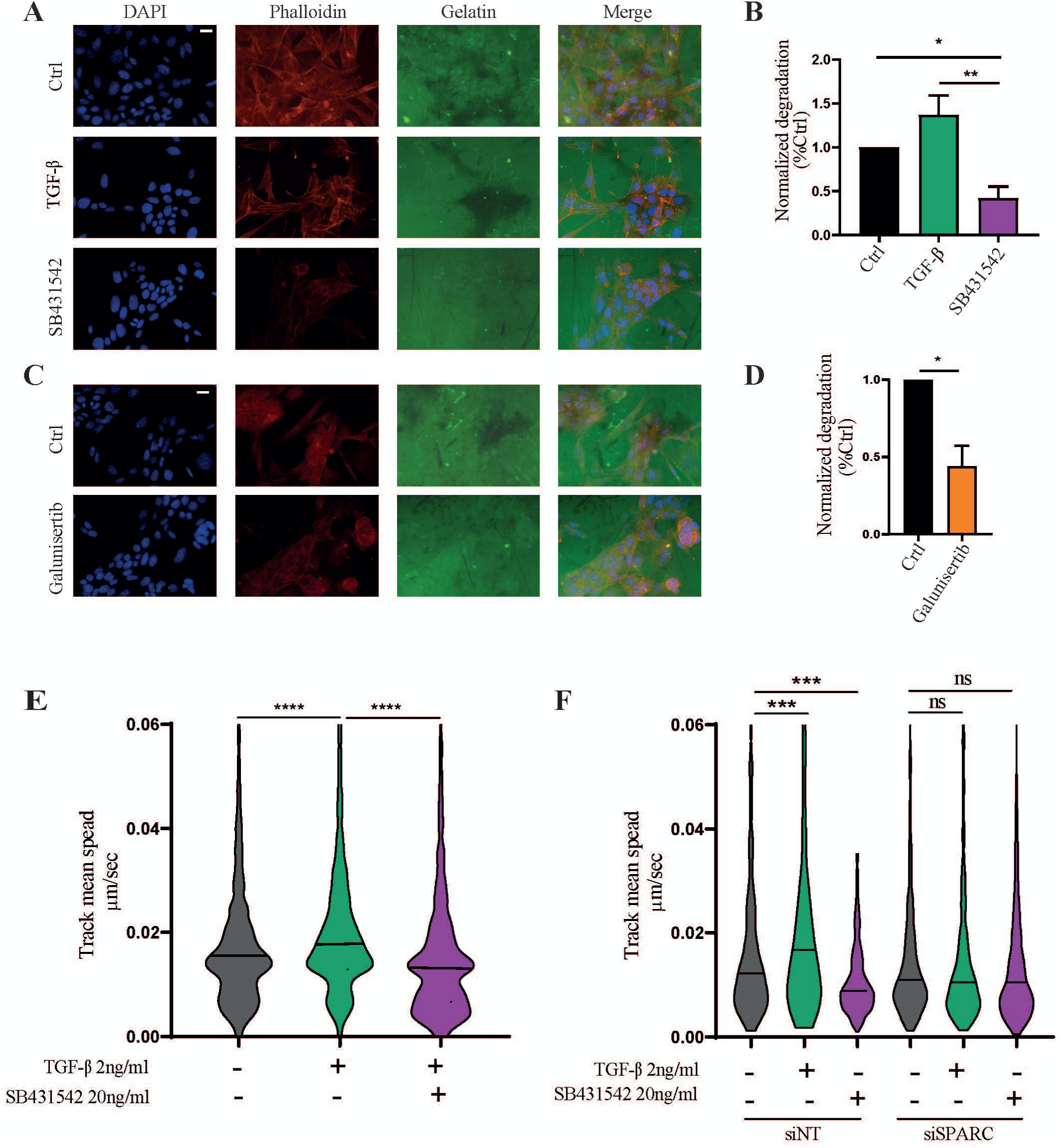
TGFBRI inhibition reduces matrix-degradative potential and motility of breast cancer cells. **A**. Representative images of the gelatin degradation assay of the murine LM38-LP cells treated or not with TGF-β (2 ng/ml) and SB431542 (20 ng/ml). Representative images of cells in fluorescent gelatin. Blue: DAPI, Green: Gelatin, Red: Phalloidin. Magnification bar: 50µm. **B**. Quantification of fluorescent gelatin degradation by total cells in LM38-LP cells, (n=4). *p<0,05, **p<0.001 using ANOVA-one way. **C.** Representative images of the gelatin degradation assay of the murine LM38-LP cells treated with Galunisertib (1 µg/ml). **D.** Quantification of fluorescent gelatin degradation by total cells in LM38-LP treated or not with Galunisertib (n=3). *p<0.05, using one tailed t-test. **E.** Single cell tracking assay in 3D collagen microdevise in LM38-LP cells treated or not with TGF-β and SB431542. Violin plot of track mean speed, *** p<0.0001, Kruskal-Wallis test. **F.** Single cell tracking assay in 2D in LM38-LP cells KO SPARC treated or not with TGF-β and SB431542. Violin plot of track mean speed, ***p<0.0001, Kruskal-Wallis test.

### Galunisertib reduces invasion in a syngeneic intraductal mouse model

To further strengthen the above-described functional association between SPARC and TGF-β pathway in an *in vivo* setting, we used the intraductal injection of LM38-LP cells in syngeneic mice (Figure supp 3A). We observed increased expression of both SPARC and TGFBRI at the invasive tumor edge (Figure supp 5E), suggesting the involvement of the TGF-β molecular pathway in the early breast cancer progression in association with SPARC. To evaluate TGF-β contribution to the *in situ*-to-invasive transition in LM38-LP intraductal tumors, tumor bearing mice were treated with Galunisertib. Based on whole-mount carmine staining of the injected mammary glands and H&E staining of tissue-sections we observed that treatment with Galunisertib decreased the proportion of invasive tumors as compared with the control untreated group (Figure 6A-C). Additionally, the IDC tumor area was significantly larger in untreated vs. Galunisertib-treated tumor xenografts indicating that the IDC component was relatively reduced (Figure 6D). Moreover, Galunisertib-treated tumors contained fewer proliferating cells, as determined by Ki67 staining, and less nuclear SMAD-4, consistent with an effective blockade of TGFBRI signalling. Furthermore, tumors from Galunisertib-treated mice showed lower SPARC and MT1-MMP intensity as compared to untreated controls (Figure 6E-F). Finally, we next examined the correlation between the expression of SPARC and TGF-β and TGFBRI using public databases. High SPARC expression was positively correlated with both TGF-β and TGFBRI expression in a basal-type human breast cancer (Figure 6G). Collectively, these data argue that the induced expression of SPARC and MT1-MMP downstream of the TGFBRI would be necessary for in situ to invasive transition in breast cancer

**Figure 6.**
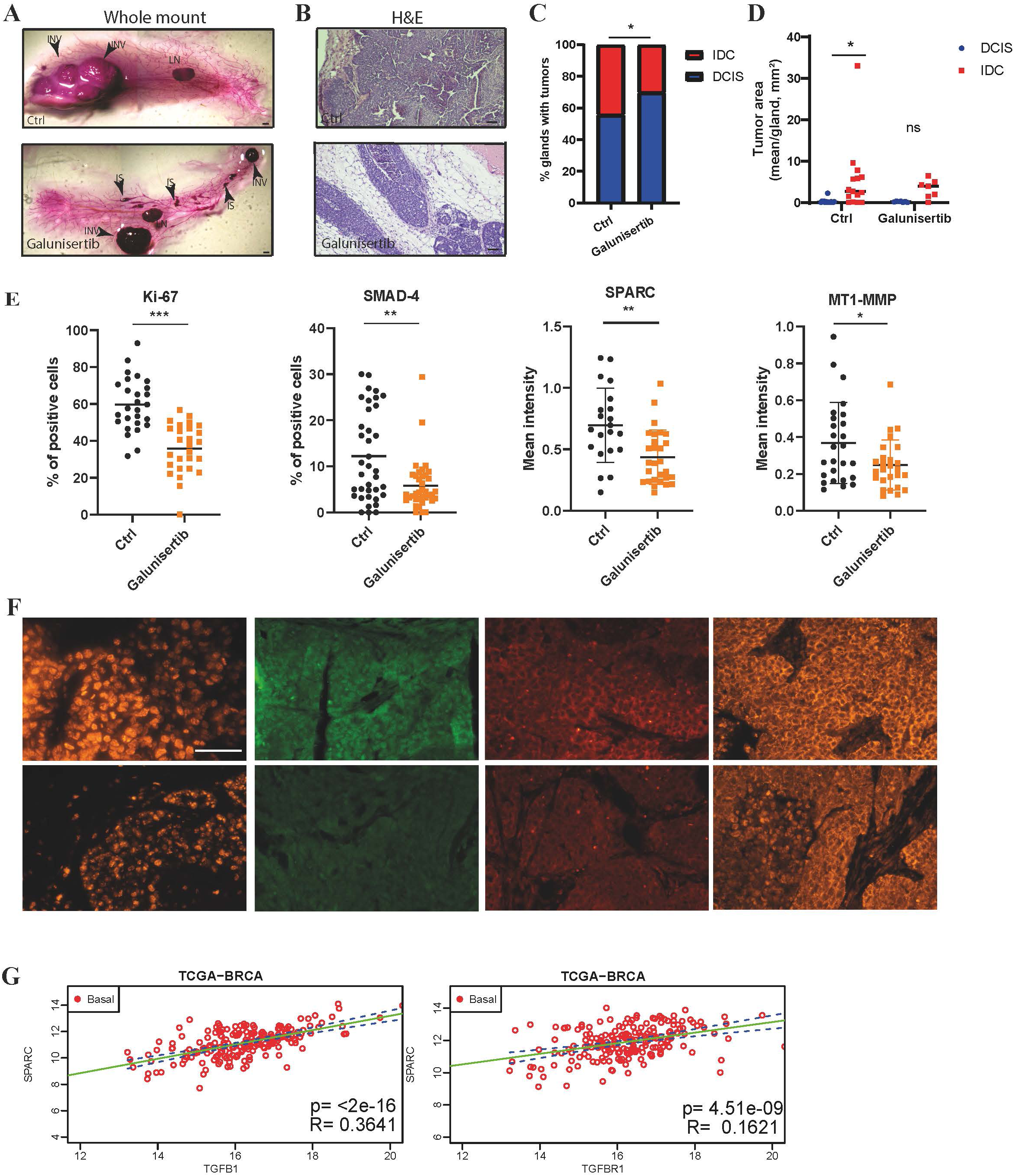
TGFBRI modulation impairs the *in situ* to invasive transition in the LM38-LP experimental model. **A.** Balb/c mice were intra-nipple injected with LM38-LP cells and treated or not with Galunisertib (60 mg/Kg). Representative images of the whole-mount carmine-stained glands analyzed 4-5 weeks after injection. IS, in situ; INV, invasive (Ctrl: n=15; Galunisertib n=18); LN, lymph node. Scale bar, 1 mm. **B.** Representative images of H&E tumors. Scale bar, 50 μm. **C.** Phenotypic analysis of intraductal xenograft tumors of LM38-LP cells control or treated with Galunisertib. Analysis was based on whole-mount and H&E staining at 4–5 weeks after intraductal injection. Phenotypic analysis of intraductal xenograft tumors of LM38-LP cells treated or not with Galunisertib. The analysis was based on whole-mount Fisher’s exact test, p= 0.0391. **D.** Tumor area per gland comparing progression status in Ctrl group and treated with Galunisertib. 2 ways ANOVA, Tukey’s multiple comparisons test, p= 0.0418. **E-F** Percentage of positive cells for Ki-67, SMAD-4, SPARC and MT1-MMP markers by Immunofluorescence analyses in tissue sections in tumors treated or not with Galunisertib with the illustrative image. **p<0.01; *** p<0.001, Mann Whitney test, (pooled tumor from n=3 glands per group). Scale bar: 50 µm. **G**. **S**catter plot displaying the relationship between SPARC and TGF-β (left panel) and TGFBRI (right panel) expression levels in breast basal cancer tumors, based on data from the TCGA breast cancer dataset. The plot includes a linear regression line (green) representing the best-fit model between the two variables, along with a 95% confidence interval for the regression predictions (blue dashed lines).

## Discussion

Understanding the molecular mechanisms that trigger the formation of invasive tumors from *in situ* lesions is of paramount importance to select appropriate treatment strategies for women diagnosed with early breast cancer. In the absence of biomarkers that can predict progression many patients receive unnecessary treatment. Experimental models can be used to address this challenge by studying the cellular mechanisms underlying the DCIS-to-IDC transition. We have previously reported an essential role of extracellular matrix-degradative and basement membrane-breaching MT1-MMP in DCIS-to-IDC transition using the human-in-mouse intraductal xenograft system of MCF10DCIS cells in immunodeficient mice^8,19,23^.

Here, using computational analysis of gene signatures obtained from MT1-MMP^high^ and MT1-MMP^low^ cells from intraductal MCF10DCIS cell tumor xeniografts showed a significant enrichment of gens involved in processes associated with cell adhesion and extracellular matrix remodelling^24^. Alignment of MT1-MMP^high^ cell and high-grade DCIS-C2 cohort^10^ gene expression profiles identified three common genes. Up-regulation of these genes in the context of DCIS-to-IDC progression both in the experimental and clinical settings prompted us to investigate these molecules as possible players in the prelude to invasive transition. Among the three candidates, SPARC, was best correlated with MT1-MMP expression. SPARC has been described pro-or anti-tumor roles in different cancers^25^, such as in advanced breast cancer patients in which low levels of SPARC protein correlated with worse prognosis as compared to tumors expressing high SPARC levels^26^. In contrast, in multivariate analysis, high SPARC expression was independently predictive for disease-free-survival in all patients^27,28^. SPARC effect appeared to depend not only on tumor molecular subtype of, but also on intratumoral cell and matrix composition ^28,29^. We thus hypothesize that SPARC play protumoral role in early breast cancer due to its involvement in DCIS-to-IDC progression.

In this work, based on IHC analysis of FFPE tumor samples from three independent cohorts totaling 230 samples and including different stages of invasion, we found that SPARC was upregulated in pre-invasive neoplastic cells compared to normal epithelium. SPARC expression level was also high in the invasive synchronous component in hormone receptor-negative tumors. In other breast cancer subsets, SPARC expression was not associated with disease progression in advanced stages. Furthermore, SPARC expression was not detected in lymph node metastases. Although strong SPARC expression in the tumor stroma was associated with shorter time to recurrence among DCIS patients^22^, in this study we centered our investigation on neoplastic epithelial cells.

Recent gene expression analysis of over 2,000 individually microdissected ductal lesions revealed that the progressive loss of basement membrane integrity, which signifies transition towards invasive carcinoma, involves two distinct epithelial-to-mesenchymal transition (EMT) events. The initial EMT event occurs early in progression, while a second event happens later, coinciding with convergence of expression profiles between ductal carcinoma in situ (DCIS) and invasive ductal carcinoma (IDC)^21^.

Overexpression of SPARC can promote cell migration and invasion and correlates with expression of mesenchymal markers in several types of cancer cells^30,31^. In addition, SPARC has been shown to increase expression or activation of a number of metalloproteinases in a fibroblastic and inflammatory context^32^ and in some cancer cells^33,34^. In this study, we refine and reinforce these observations based on public databases, IHC data in our patient cohorts and in experimental MCF10DCIS tumor xenografts. We identify SPARC-MT1-MMP cooperation as an early protumoral factor and demonstrate that MT1-MMP expression and matrix degradation is reduced in human and mouse breast cancer cells with SPARC knockdown.

Comparative analysis of differentially expressed genes between our experimental model and human high-grade DCIS, combined with co-expression and functional analysis in invasive breast cancer tumor subtypes, identified a network of genes that likely collaborate to promote tumor progression. We used STRING to examine the connectivity of the identified genes and found that TGF-β is highly interconnected with SPARC. This finding is consistent with reports indicating that SPARC acts extracellularly as a mediator of TGF-β signalling in various contexts, including fibrosis and EMT promotion in lung cancer cells^35–37^. In this study, we extend these observations to breast cancer, showing that in tumor cells of human or mouse origin, blocking TGFBRI activity reduces SPARC and MT1-MMP expression. As a consequence, the invasive proprieties of tumor cells, such as matrix degradation and motility are decreased. We speculated that DCIS-to-IDC progression is driven by TGF-β-dependent increase in SPARC and MT1-MMP levels. Secreted SPARC may interact with TGFBRI in tumor microenvironment^38^.

In vitro, treatment with SPARC did not increase p-SMAD levels in tumor cells (data not shown), suggesting that TGFBRI activation is maximal in cells in culture. On the other hand, KO SPARC cells down-modulate p-SMAD2/3. One possible scenario is that basal TGF-β activity induces SPARC expression as part of the first wave of EMT, while secreted SPARC amplifies TGFBRI signaling to induce MT1-MMP expression, participating in the second EMT wave and to DCIS-to-IDC transition. This leads us to hypothesize that basal levels of SPARC, when secreted, activates signaling molecular pathway trigged by the TGFBRI such as MT1-MMP expression, one of the target genes of this pathway.

LM38-LP, a triple negative mouse cell line of breast cancer, was positive for SPARC and MT1-MMP. In this work we described for the first time that this cellular model is able to develop *in situ* tumors after intraductal injection in syngeneic mice. These intraductal tumors spontaneously progress into invasive ones, which makes this model an excellent tool for studying molecular factors involved in early breast cancer progression events. During progression, as soon as microinvasive cell foci appear, recruitment of inflammatory cells is observed (data not shown). Tumor-bearing mice treated with Galunisertib reduced the proportion and tumor area of invasive foci. The proinflammatory role of TGF-β is well known during tumor progression^39^ and although we cannot rule out that Galunisertib is acting by blocking the immune system, the tumor cells treated with the inhibitor present lower SMAD-4 positive cells compared to Controls. Thus, TGF-β signaling pathway is being blocked by the inhibitor in tumor cells. Similar to what we observed *in vitro*, SPARC and MT1-MMP expression is in consequence also reduced in Galunisertib treated tumours. All together, these findings lead us to propose that a mechanism that involves activated TGF-β pathway, induces sustained expression levels of SPARC and MT1-MMP, which are responsible for triggering pro-invasive mechanisms responsible for early transition in triple-negative breast cancer. Lately, efforts have been made to develop specific tools that allow better understanding of the molecular signatures accompanying mechanisms that lead to the progression of carcinoma *in situ*^40^. In this work, we identify SPARC as an essential gene included in a molecular pathway involved during the DCIS-to-IDC transition, and we propose that the TGFBRI/SPARC/MT1-MMP axis may offer therapeutic targets to improve management of patients with *in situ* tumors.

## Acknowledgements

The authors acknowledge the Breast Cancer Study Group and the Incentive and Cooperative Research Program “Breast cancer: Cell Invasion and Motility” of Institut Curie and all the staff of the Department of Pathology of the Diagnostic Area of the Angel Roffo Institute, specially Dr Fernando Carrizo, Fernanda Rocca and Gisela Lopez for their continued support. We also thank the patients from both institutions for the breast tumor samples. Dr Cecilia Perez Piñero and M. Sol Rodriguez for IHQ of PR and ER assistance. Authors also thank Dr Rebbeck for generously providing the timeline data documenting progression from normal breast tissue to invasive breast cancer. We also thank A. V. Failla and the UKE Microscopy Imaging Facility (UMIF) under the DFG Research Infrastructure Portal: RI−00489.

## Author contribution

M.S. performed in vitro and vivo experiments, analysed the data and participated in the article conceptualization. M.P.C. contributed for the collection of the samples at the Institut, L.M. performed IHQ staing. M.P.C., C.L. and L.M. analysed clinical samples. E.L. and M.A performed the RNAseq analysis and analysed public transcriptome data set. N.P.C. assisted in vivo experiments. N.R.P, T.L.L. and K.P. performed in vitro experiments. AV-S selected and classified patients from PICBIM. E.R.B. supervised patient’s analysis. L.C and I.B. provided and analyzed clinical sample data. M.B., P.C., A.M.E. and P.S. assisted with resources, review and editing. C.L. supervised the project, designed experiments and wrote the manuscript.

**Figure supplementary 1. SPARC expression in advanced stages. A.** SPARC levels using the H-score method in the adjacent peritumoral tissues, invasive breast carcinomas and in lymph node metastasis (LNM) in Roffo’s cohort. EPI vs IDC** p=0.0031, IDC vs LNM* p= 0.0450 Kruskal-Wallis test. **B.** Proportion of negative or positive tumors for SPARC EPI vs IDC, **p=0.0017, IDC vs LNM *p= 0.03, X^2^ test, two-side. **C**. Representative SPARC IHC staining in lymph node metastasis. **D**. Metastasis-free survival (MFS) curves for IDC tumor patients with low and high SPARC expression by IHQ.

**Figure supplementary 2. SPARC and MT1-MMP expression in cell line collection A**. Expression levels of MT1-MMP (left panel) and SPARC (right panel) in a collection of breast cancer cell lines from the GSE48213 dataset, categorized by Luminal (blue) and Basal (red) subtypes. **B.** Metastasis-free survival (MFS) curves for breast tumor patients with low and high MT1-MMP (left panel, p=0.014) SPARC mRNA expression (middle panel, NS) and the combination of both markers (NS) mRNA expression as compared with normal breast tissue. The 458 breast tumors were divided into two groups with low (<3) and high (>3) for both markers.

**Figure supplementary 3. A new syngenic intraductal model. A.** LM38-LP intraductal model characterization. Balb/c mice were intra-nipple injected with LM38-LP cells in the foth mammary glands. Whole-mount and H&E images of *in situ*, microinvasive and invasive stage of tumors, at 3, 4 and 5 weeks respectively after intraductal inoculation. Scale bar: 50 µm. **B.** Representative ER, PR and HER2/neu IHC staining in invasive tumors after intraductal injection of LM38-LP. Insets corresponds to positive corresponding control: normal epithelial duct (PR and ER) and Positive HER2/neu breast tumor. Scale bar: 100 µm. **C.** Immunofluorescence analysis of α-SMA (red) and DAPI (blue) in *in situ* (upper panel) or invasive (lower panel). Scale bar: 50 µm.

**Figure supplementary 4. Knockdown of SPARC by RNAi did not affect cell proliferation or morphology. A.** Immunofluorescence staining for MT1-MMP (red) and SPARC (red) expression in LM38-LP and MCF10DCIS wild type cells. Scale bar: 10 µm. **B.** Contrast face images illustrating morphology of LM38-LP and MCF10DCIS cells transiently silenced or not for SPARC. Scale bar: 50 µm. **C.** Quantification of viable Trypan blue LM38-LP and MCF10DCIS cells transiently silenced or not for SPARC. **D**. Immunofluorescence staining for MT1-MMP (red) and SPARC (red) expression in MCF10DCIS cells transiently silenced or not for SPARC. Scale bar 10 um. **E**. Quantification of fluorescent signal for MT1-MMP and SPARC expression in MCF10DCIS cells transiently silenced or not for SPARC. *** p<0.0001 and *p<0.05 using two tailed t-test.

**Figure supplementary 5. Identification of co-expressed genes. A**. Euler diagram representing the number of genes co-expressed with each of the 4 genes and the overlap between them. **B**. Number of genes co-expressed by SPARC and MT1-MMP in the 6 groups of samples considered. **C.** mRNA levels of SPARC (left panel) and MT1-MMP (right panel) in MCF10DCIS cells after SB431542 treatment (20 ng/ml). Values corresponding to means ± SEM, n = 3. t-test, **p<0.001. **D.** Single cell tracking assay in 3D collagen microdevice in MCF10DCIS cells treated or not with TGF-β and SB431542. Violin plot of track mean speed, ***p<0.0001, Kruskal-Wallis test. **E.** Immunofluoresce illustrating SPARC and TGBRI expression in invasive tumors after intraductal injection of LM38-LP. Scale bar: 10 µm.

