## Supplemental Figure 1-5 for "SPARC is a new driver of early breast tumor progression via TGF-β -dependent mechanism"

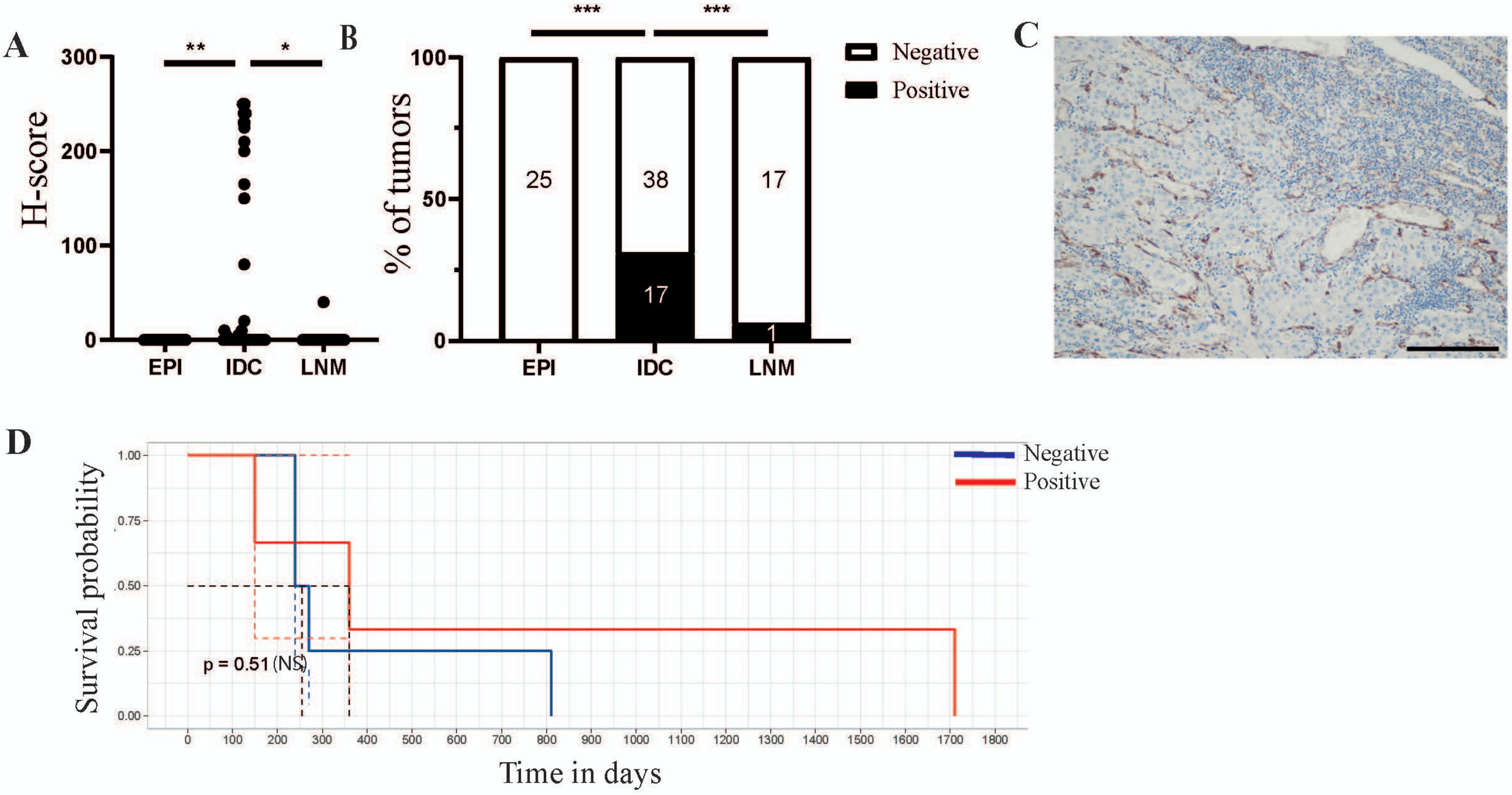

Supplementary Figure 1

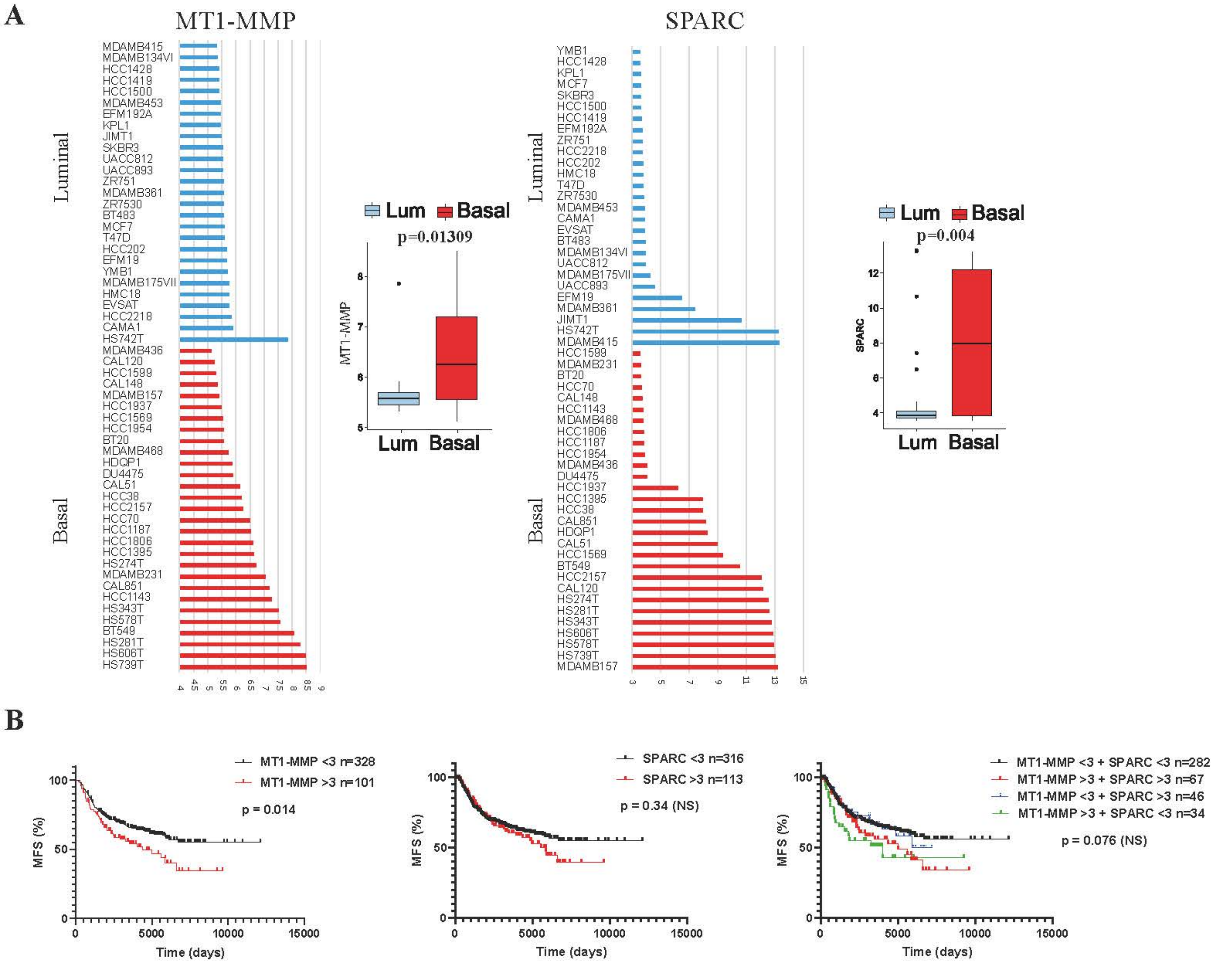

Supplementary Figure 2

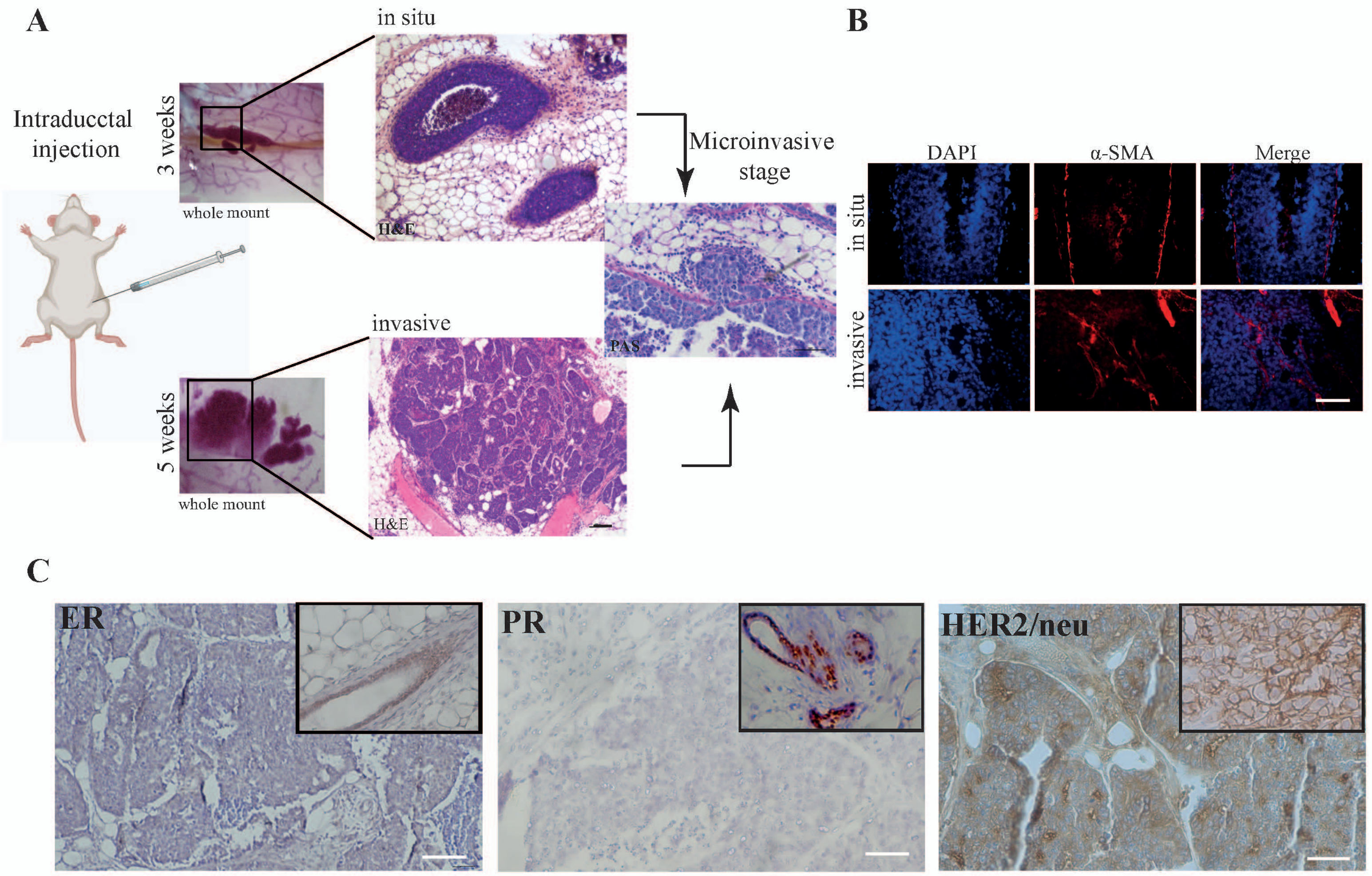

Supplementary Figure 3

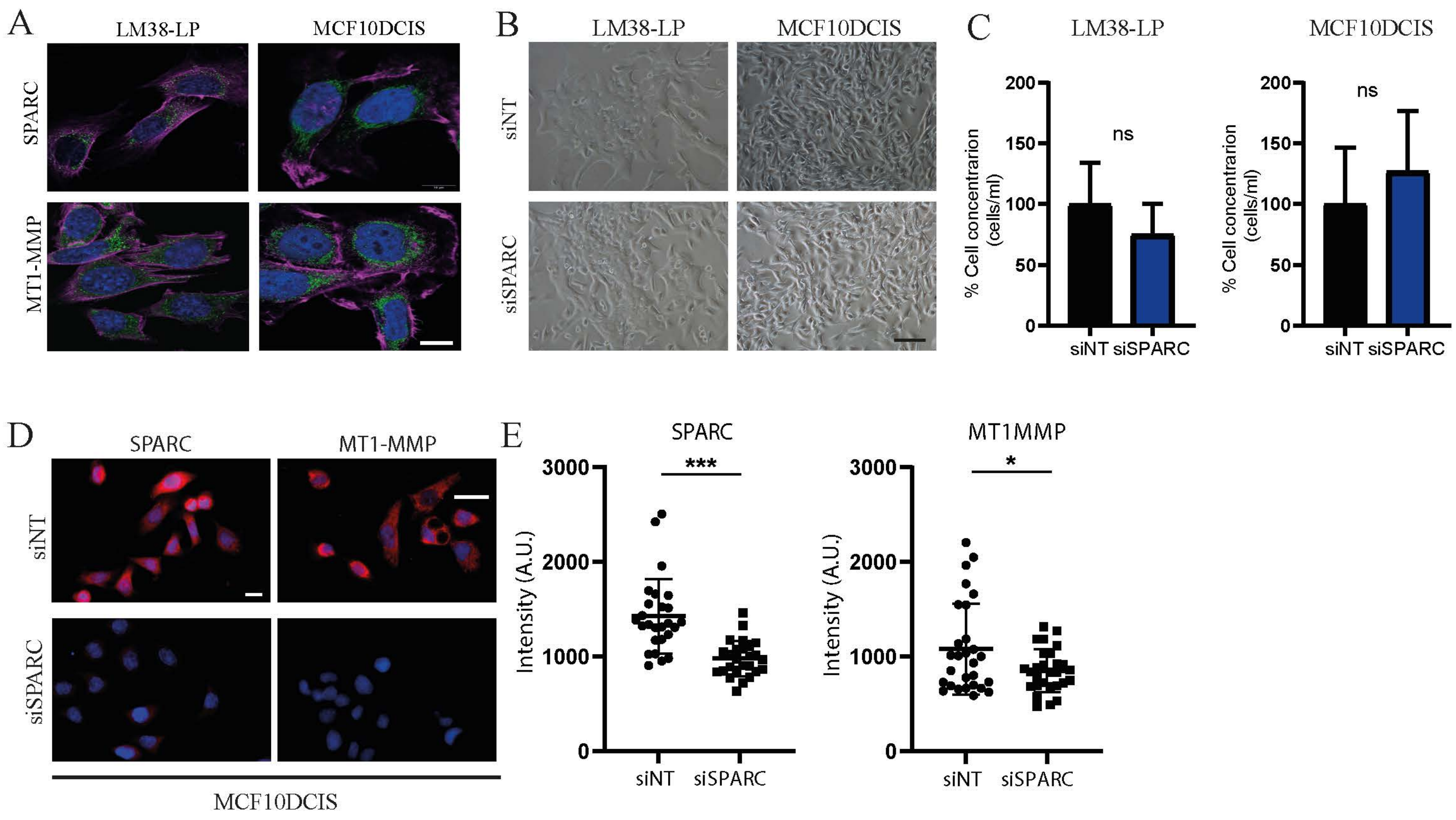

Supplementary Figure 4

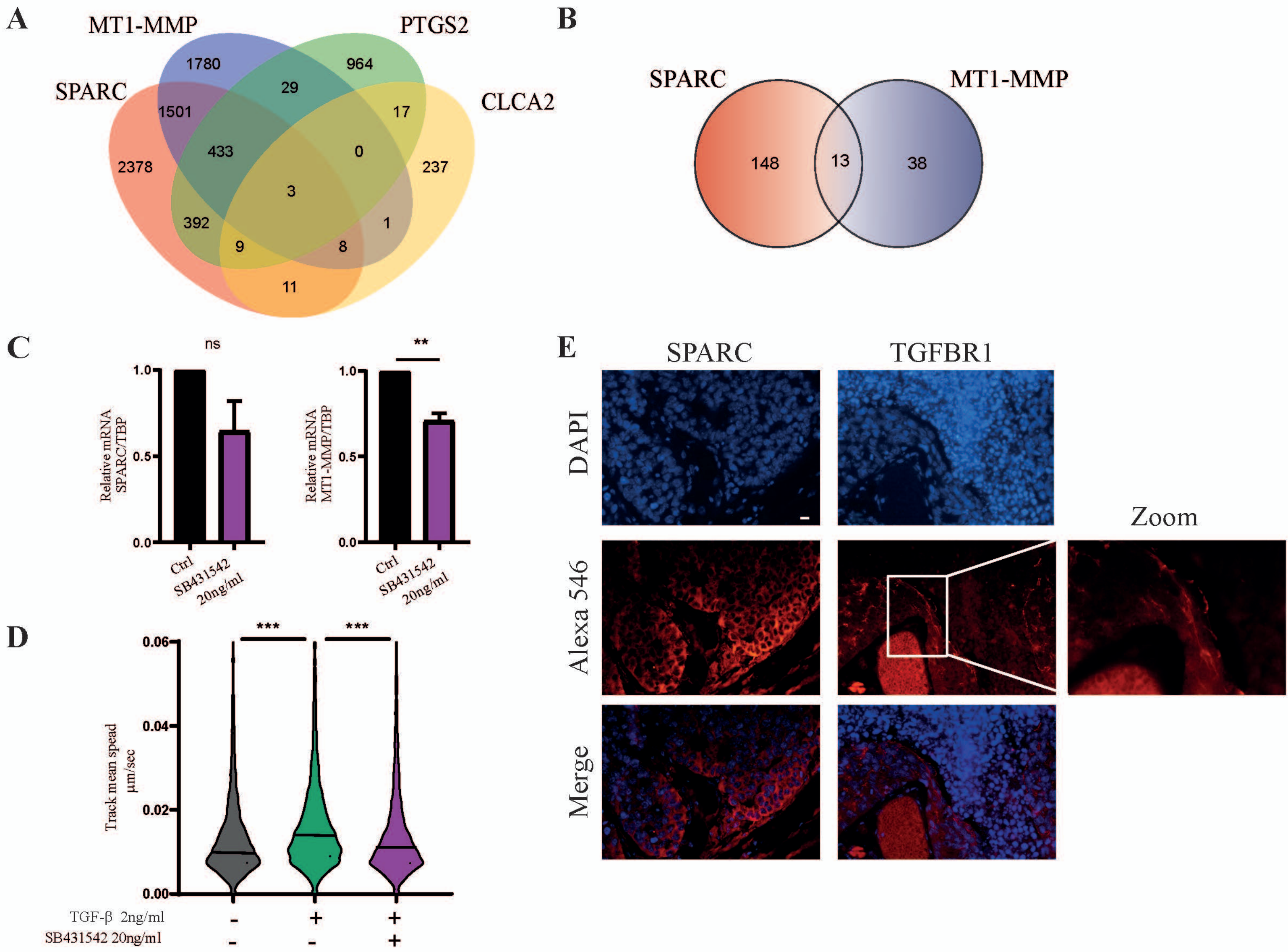

Supplementary Figure 5

| FEATURES COHORT I | Invasive carcinoma (n=116) |
|---|---|
| *Age (years)* | |
| ≤ 50 | 46 (39.7%) |
| > 50 | 69 (59.4%) |
| Unknown | 1 (0.9%) |
| *Histological grade* | |
| I | 12 (10.3%) |
| II | 35 (30.2%) |
| III | 68 (58.6%) |
| Unknown | 1 (0.9) |
| *In situ component* | |
| present | 78 (67.3%) |
| absent | 34 (29.3%) |
| NA | 4 (3.4%) |
| *Histological subtype* | |
| Ductal carcinoma | 116 (100%) |
| Others | 0 (0%) |
| *Tumour size (cm)* | |
| Tx | 1 (0.9%) |
| T1 (<2) | 81 (69.8%) |
| T2 (2 - 5) | 33 (28.4%) |
| T3 (>5) | 1 (0.9%) |
| *N stage* | |
| NX | 1 (0.9) |
| N0 | 64 (55.2%) |
| N1 | 39 (33.6%) |
| N2 | 8 (6.9%) |
| N3 | 4 (3.4%) |
| *ER* | |
| Positive | 38 (32.7%) |
| Negative | 78 (67.3%) |

| | |
|---|---|
| Unknown | 0 (0%) |
| *PR* | |
| Positive | 35 (30.2%) |
| Negative | 81 (69.8%) |
| Unknown | 0 (0%) |
| *HER2* | |
| Positive | 19 (16.4%) |
| Negative | 97 (83.6%) |
| Unknown | 0 (0%) |
| *Ki67* | |
| Positive (≥20%) | 95 (81.9%) |
| Negative (<20%) | 19 (16.4%) |
| Unknown | 2 (1.7%) |
| *Molecular subtype* | |
| TNBC | 61 (52.6%) |
| HER2 | 18 (15.5%) |
| Luminal A | 20 (17.2%) |
| Luminal B | 16 (13.8%) |
| Luminal B / HER2 | 1 (0.9%) |

| FEATURES COHORT II | Invasive carcinoma (n=42) | In situ Carcinoma (n=16) |
|---|---|---|
| *Age (years)* | | |
| ≤ 50 | 14 (33.4%) | 6 (37.5%) |
| > 50 | 28 (66.6%) | 10 (62.5%) |
| *Histological grade (invasive tumors)* | | |
| I | 12 (28.6%) | x |
| II | 18 (42.8%) | x |
| III | 12 (28.6%) | x |
| *In situ component* | | |
| present | 34 (81%) | x |
| absent | 8 (19%) | x |
| *Nuclear grade (DCIS)* | | |
| High | x | 15 (94%) |
| Non high | x | 1 (6%) |
| *Histological subtype* | | |
| Ductal carcinoma | 38 (90.5%) | x |
| Others | 4 (9.5%) | x |
| *Architectural pattern (DCIS)* | | |
| Solid | x | 10 (62.5%) |
| Cribriform | x | 4 (25,0%) |
| Papillary | x | 1 (6.25%) |
| Other | x | 1 (6.25%) |
| *Tumour size (cm)* | | |
| Tis | x | 16 (100) |
| T1 (<2) | 22 (52.4%) | x |
| T2 (2 - 5) | 16 (38.1%) | x |
| T3 (>5) | 1 (2.4%) | x |
| Unknown | 3 (7.1%) | x |
| *N stage* | | |
| N0 | 33 (78,6%) | x |

| | | |
|---|---|---|
| N1 | 6 (14.3%) | x |
| N2 | 2 (4.7%) | x |
| N3 | 1 (2,4%) | x |
| *ER* | | |
| Positive | 32 (76.2%) | 13 (81.2%) |
| Negative | 9 (21.4%) | 2 (12.5%) |
| Unknown | 1 (2.4%) | 1 (6.3%) |
| *PR* | | |
| Positive | 28 (66.7%) | 12 (75.0%) |
| Negative | 13 (30.9%) | 3 (18.7%) |
| Unknown | 1 (2.4%) | 1 (6.3%) |
| *HER2* | | |
| Positive | 5 (11.9%) | 15 (40,6%) |
| Negative | 36 (85.7%) | 22 (59,4%) |
| Unknown | 1 (2.4%) | |
| *Ki67* | | |
| Positive (≥20%) | 24 (57.1%) | x |
| Negative (<20%) | 17 (40.5%) | x |
| Unknown | 1 (2.4%) | |
| *Molecular subtype* | | |
| TNBC | 6 (14,3%) | |
| HER2 | 3 (7.2%) | |
| Luminal A | 16 (38,1%) | |
| Luminal B | 15 (35,7%) | |
| Luminal B / HER2 | 2 (4.7%) | |

| FEATURES COHORT III | Invasive carcinoma (n=57) |
|---|---|
| *Age (years)* | |
| ≤ 50 | 21 (36.8%) |
| > 50 | 5 (61.4%) |
| Unknown | 1 (1.8%) |
| *Histological grade (invasive tumors)* | |
| I | 8 (14.0%) |
| II | 23 (40.4%) |
| III | 20 (35.1%) |
| Unknown | 6 (10.5%) |
| *Histological subtype* | |
| Ductal carcinoma | 47 (82.5%) |
| Others | 8 (14.0%) |
| Unknown | 2 (3.5%) |
| *Tumour size (cm)* | |
| T1 (<2) | 29 (50.9%) |
| T2 (2 - 5) | 12 (21.0%) |
| T3 (>5) | 4 (7.0%) |
| T4 | 4 (7.0%) |
| Unknown | 8 (14.1%) |
| *N stage* | |
| N0 | 21 (36.8%) |
| N1 | 18 (31.6%) |
| N2 | 6 (10.5%) |
| N3 | 5 (8.8%) |
| Unknown | 7 (12.3) |
| *ER* | |
| Positive | 32 (56.1%) |
| Negative | 23 (40.3%) |
| Unknown | 2 (3.6%) |

| | |
|---|---|
| *PR* | |
| Positive | 24 (42.1%) |
| Negative | 31 (54.3%) |
| Unknown | 2 (3.6%) |
| *HER2* | |
| Positive | 15 (26.3%) |
| Negative | 39 (68.4%) |
| Unknown | 3 (5.3%) |
| *Ki67* | |
| Positive (≥20%) | 39 (68.4%) |
| Negative (<20%) | 15 (26.3%) |
| Unknown | 3 (5.3%) |
| *Molecular subtype* | |
| TNBC | 16 (28,1%) |
| HER2 | 8 (14.0%) |
| Luminal A | 12 (21.0%) |
| Luminal B | 11 (19.3%) |
| Luminal B / HER2 | 7 (12.3%) |
| Unknown | 3 (5.3%) |
